## Supplemental Files for "Hippocampal neurons code individual episodic memories in humans"

**This PDF file includes:**

Figures S1 to S7

Tables S1 and S2

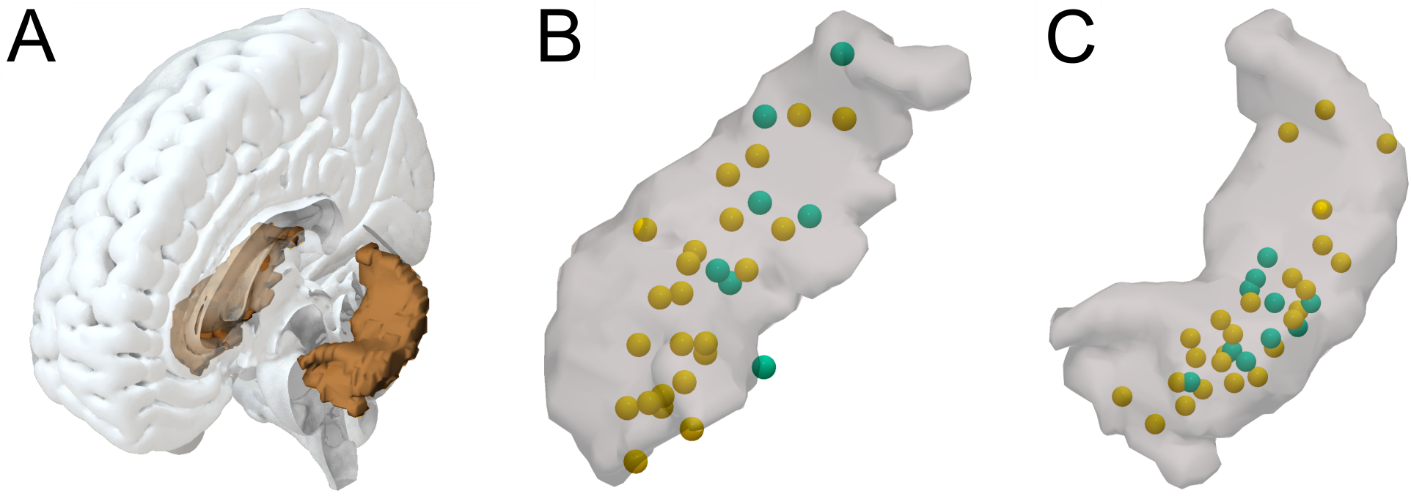
Figure S1. Visualisation of the hippocampus and electrode positions for experiment 1 and experiment 2.

(A) Outline of the hippocampus within a whole brain mesh.
(B). Normalized right hippocampus. The yellow spheres represent the estimated position of microwire bundles that contain ESNs. The green spheres represent the estimated position of microwire bundles that do not contain ESNs. Only bundles where single-unit activity was recorded are shown.
(C) Same as (B), but for the left hippocampus.

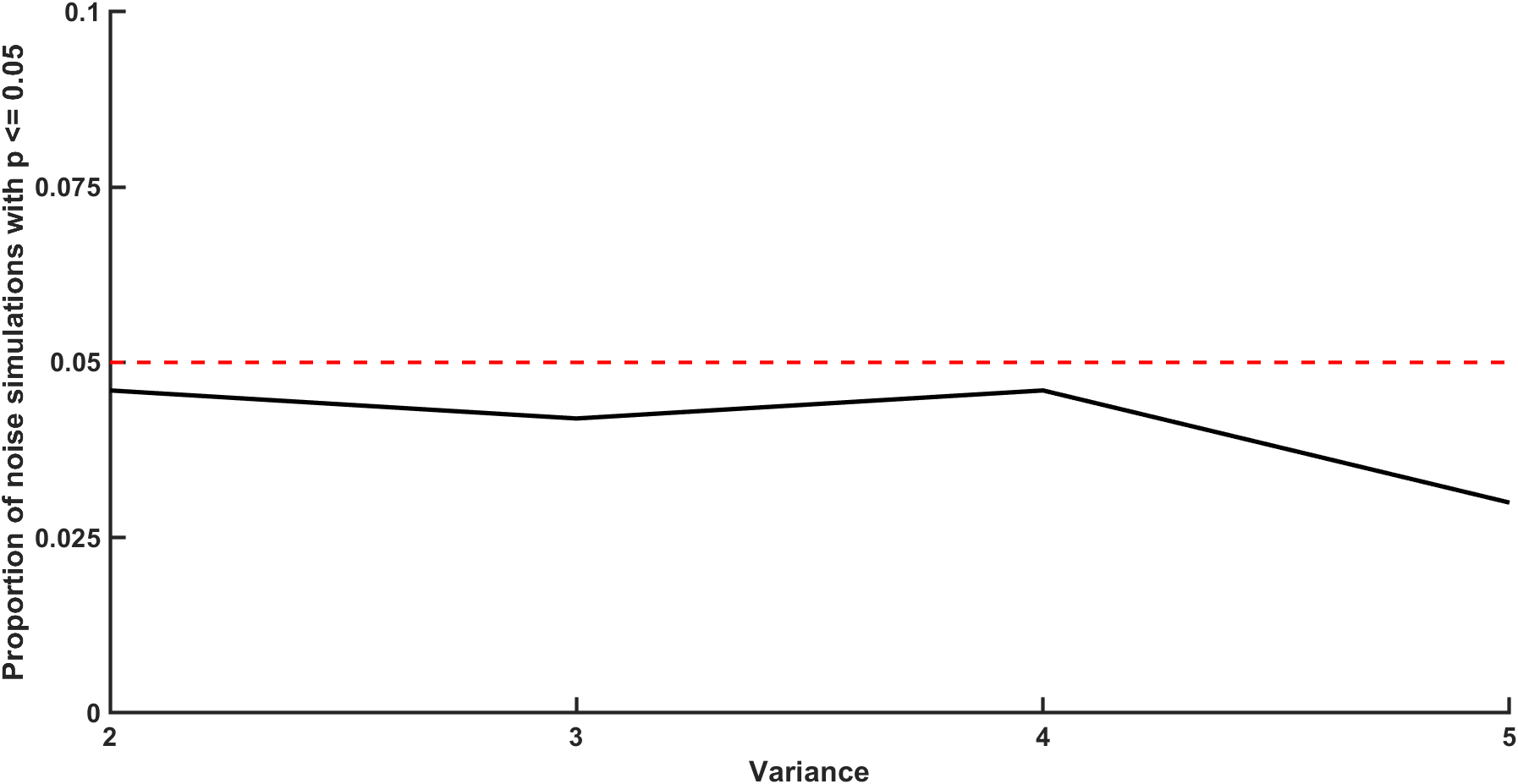

Figure S2. Simulation of ESN identification.

We created a simulation using random pseudo spike rates to determine whether our ESN analysis pipeline contains a bias towards significant results over multiple levels of variance (x-Axis). For each level of variance, we repeated this step 1,000 times and calculated the proportion of iterations that yield a significant result (y-axis; p <= 0.05). The dotted red line represents the 5%-level, and the straight black line represents the results from the stimulation.

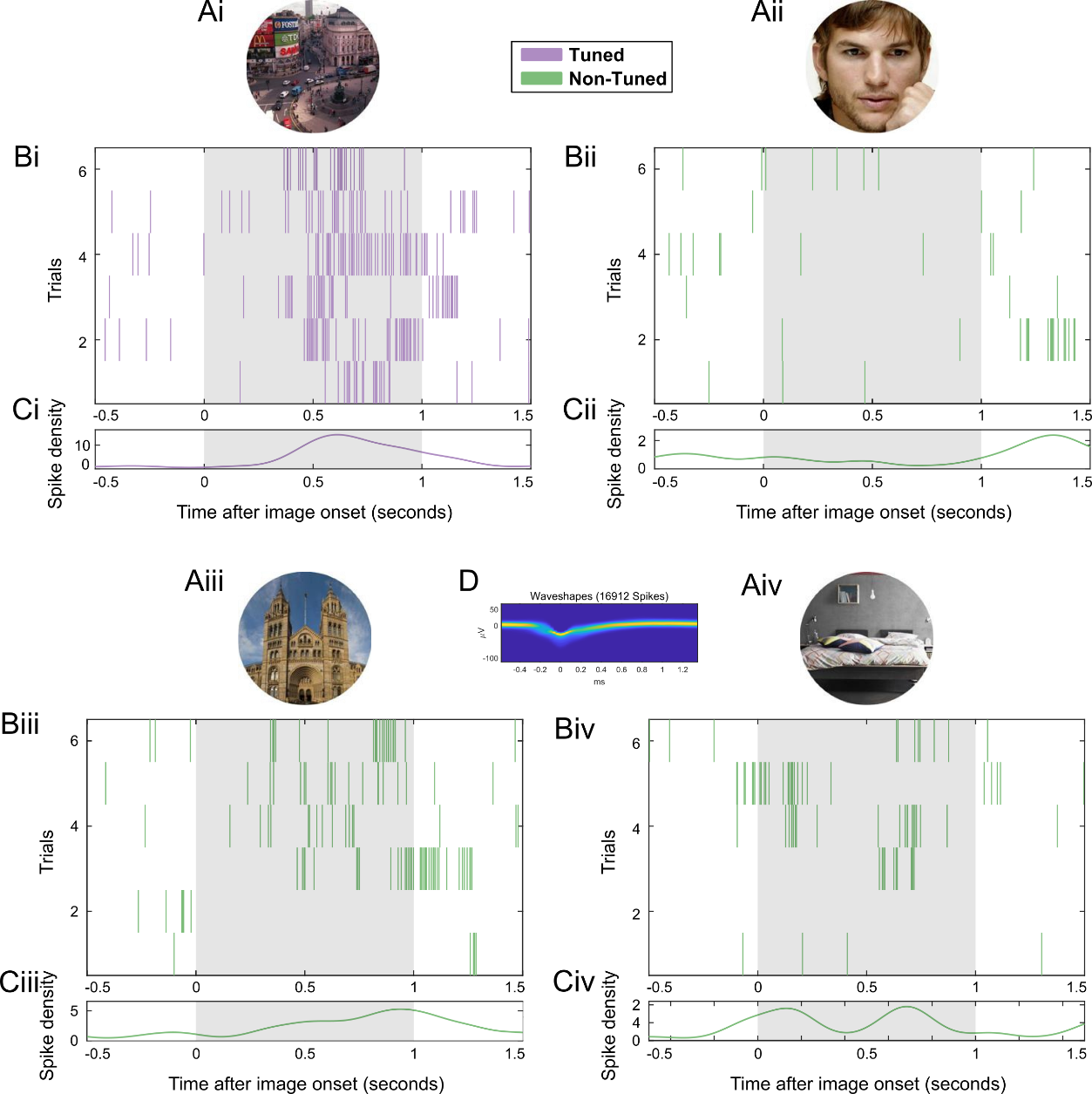

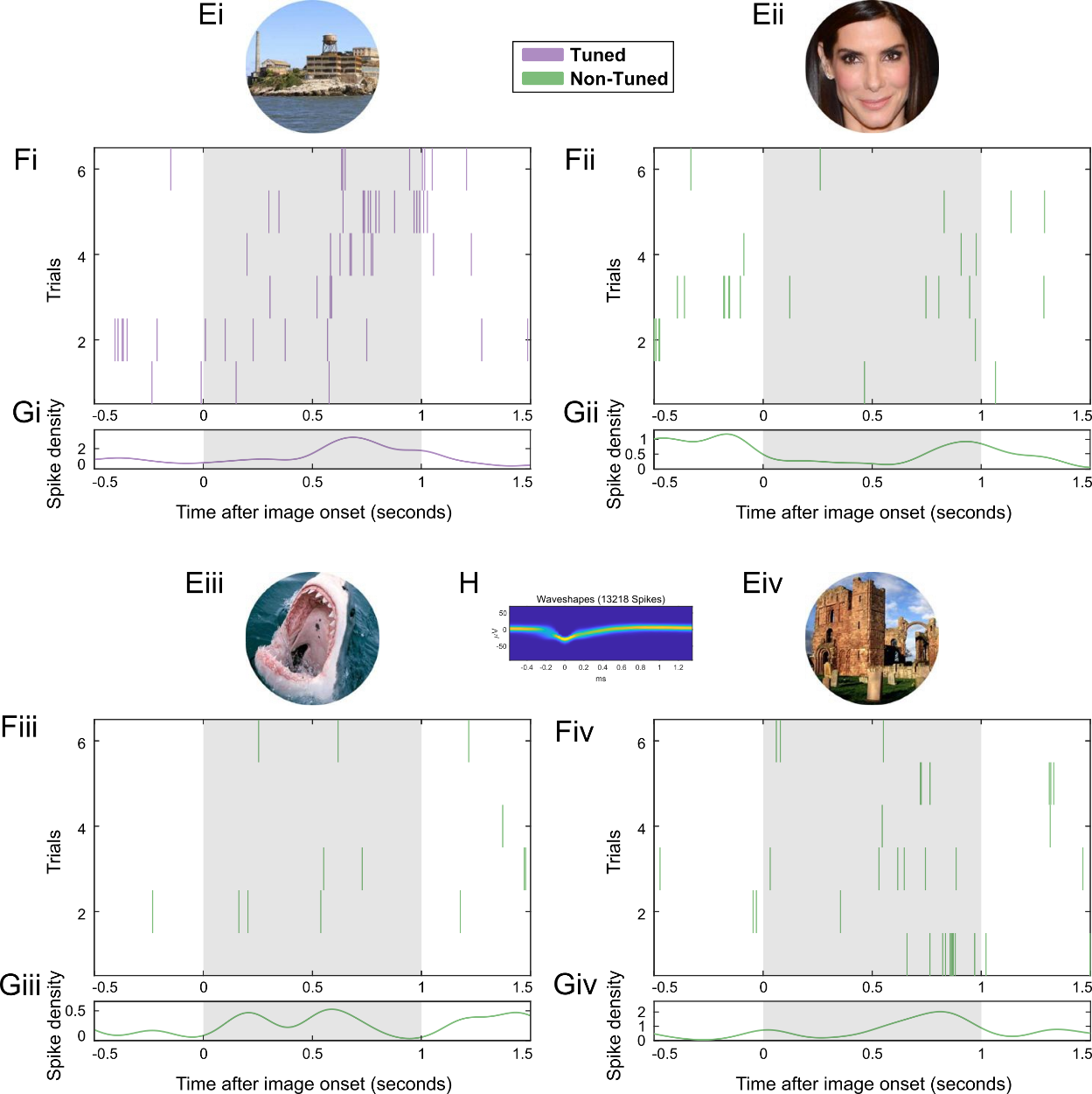

Figure S3. Firing patterns for two example putative Concept Neurons that were identified using a visual tuning task in experiment 2.

(A) Each during the memory task previously shown image (either an animal, a face or a place) is shown six times during the visual tuning task.

(B) Spike raster plot. Each line indicates a spike. On the x-axis is time (locked to image presentation) and on the y-axis are the six trials during which the above image is shown on the screen. Color-coded in purple for tuned images and green for non-tuned images. The grey area indicates the activation period that is considered for identifying Concept Neurons.

(C) Spike density plot (mean instantaneous firing rate over all six trials).

(D) 2D histogram of the waveshape of that particular unit (Niediek et al., 2016).

(E-H) same as (A-D) but for a different example ESN.

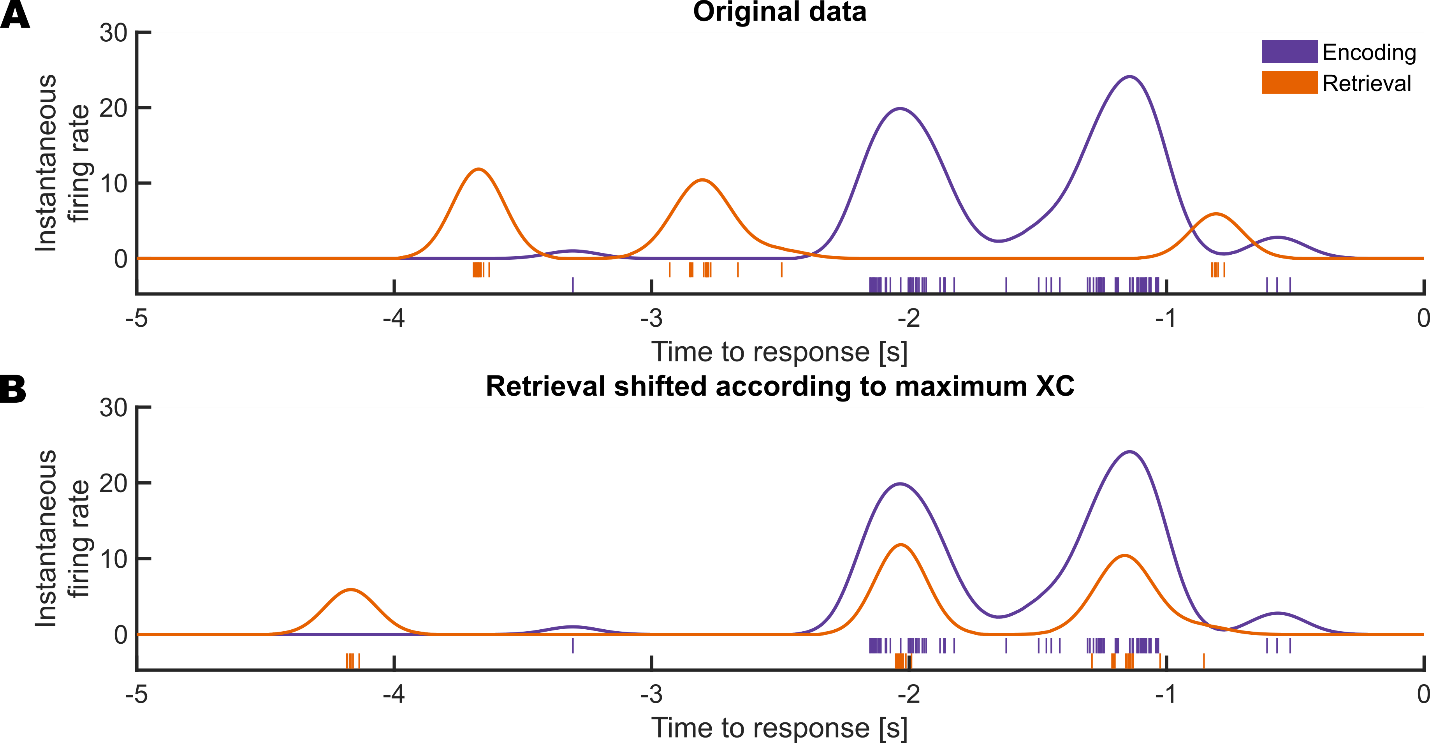

Figure S4. Example of a trial reinstated by an example temporal Episode Specific Neuron (tESN).

(A) The original instantaneous firing rate during encoding (blue) and retrieval (orange) 5000ms before the response. Each vertical line represents one action potential.

(B) Same as (A), but with the retrieval firing rate shifted according to the peak in the cross-correlation.

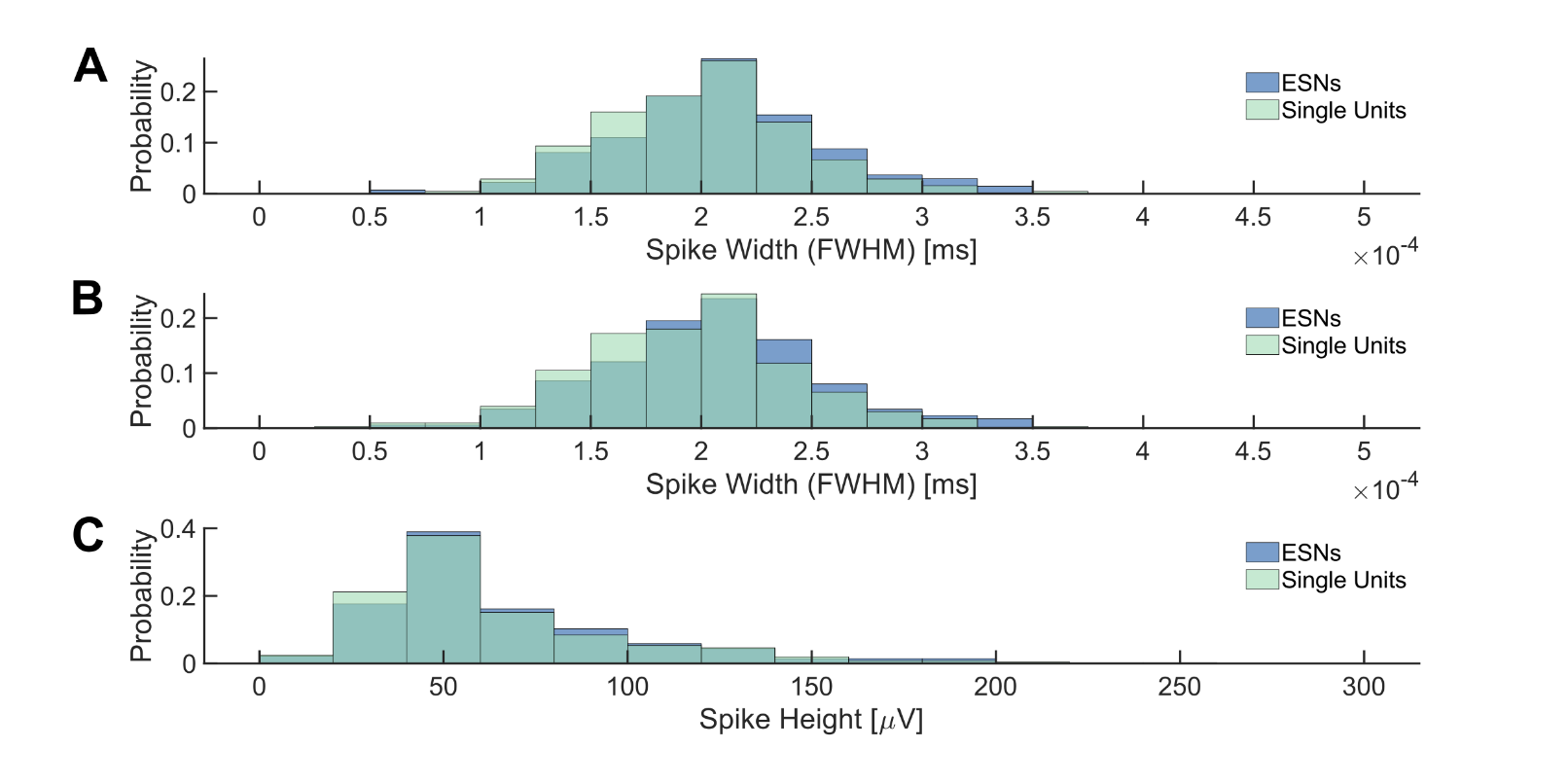

Figure S5.

(A) Distribution of spike widths for neurons of the first experiment.

(B) Same as (A) but for neurons of the first and second experiment combined.
(C) distribution of spike height of ESNs (purple) and single units (green).

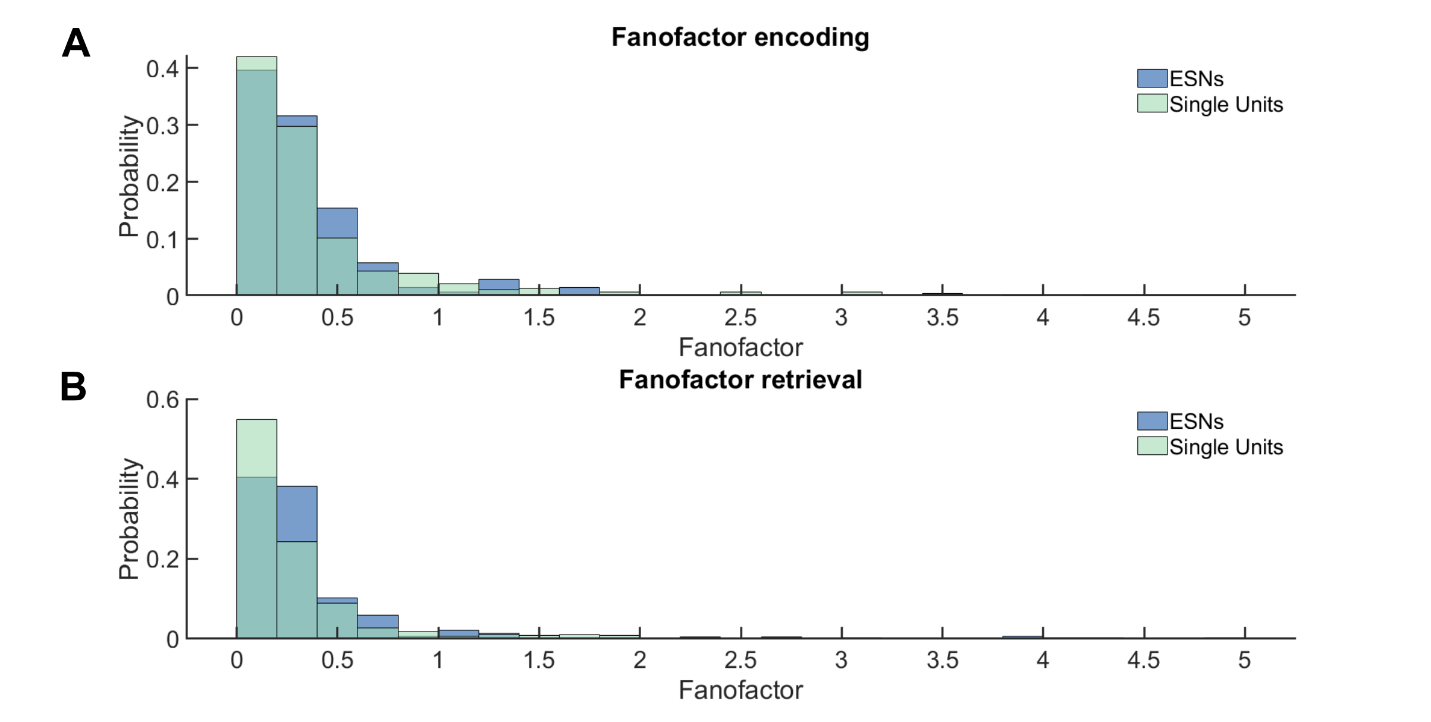

Figure S6.
(A) Distribution of Fano factors of ESNs (purple) and single units (green) during
encoding and during
(B) retrieval.

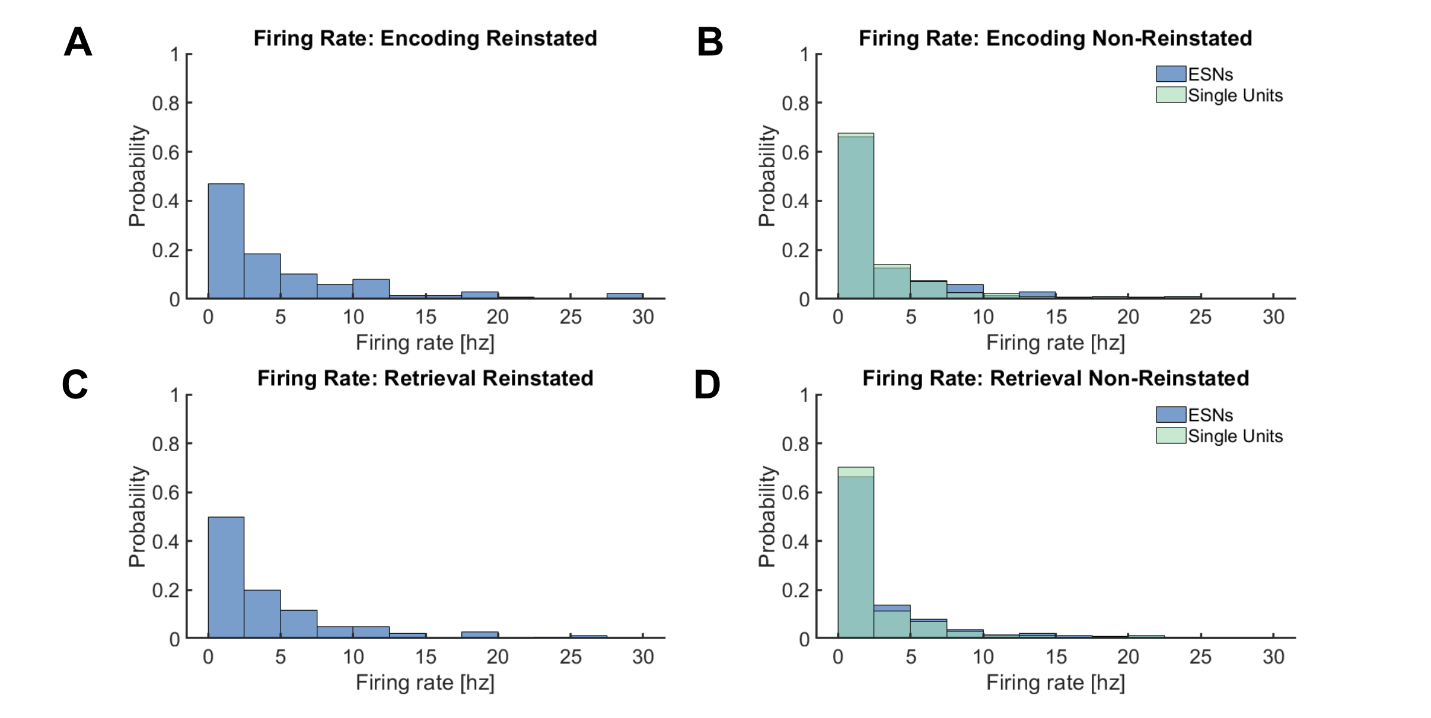

Figure S7.

Firing rate in hertz of ESNs (purple) during
(A) encoding and
(B) retrieval of reinstated episodes. Firing rate of ESNs and other single units (green) during
(C) encoding and
(D) retrieval of non-reinstated episodes.

Table S1.

| **Patient ID** | **Number of sessions** | **Number of bundles** | **Number of bundles in hippocampus** | **Trial number**^a^ | | **Hits**^a^ | **Hipp.**  **bundles with SUs^a,b^** | | **Number of hipp. SUs^a,b^** | **Number of ESNs^a,c^** | **SOZ^d^** |
| --- | --- | --- | --- | --- | --- | --- | --- | --- | --- | --- | --- |
| 0002 | 7 | 6 | 6 | 49.4 (0.4) | 43.6 (1.3) | | | 2.6 (0.3) | 12.3 (2.9) | 2.7 (0.64) | hip. (bi) |
| 0004 | 3 | 4 | 3 | 49 (0) | 33.7 (0.7) | | | 2 (0) | 7.3 (2) | 1 (0.58) | ex. (left) |
| 0005 | 4 | 6 | 6 | 49 (0) | 41.5 (2.7) | | | 3.3 (0.3) | 10.3 (1.5) | 3.8 (0.63) | ex. (right) |
| 0007 | 3 | 4 | 4 | 94.3 (1.7) | 86.7 (4.9) | | | 4 (0) | 16.7 (4.9) | 8.7 (2.6) | hip. (right) |
| 0008 | 4 | 6 | 4 | 68.8 (2.3) | 39.3 (3.9) | | | 1.8 (0.3) | 8.3 (1.6) | 1.5 (0.29) | hip. (bi) |
| 0009 | 3 | 8 | 5 | 84.3 (7.3) | 56.3 (7.3) | | | 4 (0) | 17 (1.2) | 2.7 (1.2) | hip. + ex. (left) |
| 0012 | 4 | 8 | 6 | 53.8 (8.1) | 32.5 (6.9) | | | 4 (0.4) | 21.5 (4.1) | 4.5 (0.87) | hip. (left) |
| 0013 | 6 | 5 | 5 | 76.2 (6.3) | 45.3 (8.6) | | | 2.8 (0.3) | 17.2 (1.6) | 3.7 (1.3) | ex. (left) |
| 1003 | 4 | 2 | 1 | 52.5 (2.4) | 46.3 (4.9) | | | 1 (0) | 6.3 (0.5) | 1.5 (0.87) | ex. (left) |
| 1004 | 2 | 2 | 2 | 84.5 (11.5) | 75 (11) | | | 1 (0) | 1 (0) | 1 (0) | ex. (left) |
| 1005 | 4 | 2 | 2 | 73 (13.9) | 33.3 (7) | | | 1.5 (0.3) | 3.5 (1) | 0.5 (0.29) | hip (right) |
| 1007 | 5 | 2 | 1 | 49.6 (8.5) | 31 (6.3) | | | 1 (0) | 4.4 (0.5) | 0.8 (0.37) | ex. (left) |
| 1008 | 3 | 1 | 1 | 85 (9.5) | 22.3 (6.8) | | | 1 (0) | 1.7 (0.7) | 0 (0) | ex. (left) |
| 1009 | 2 | 2 | 1 | 82.5 (13.5) | 67.5 (12.5) | | | 1 (0) | 2.5 (0.5) | 0 (0) | ex. (bi) |
| 1011 | 2 | 2 | 2 | 51 (10) | 38 (5) | | | 1.5 (0.5) | 8.5 (0.5) | 1.5 (1.5) | hip. (left) |
| 1012 | 3 | 2 | 2 | 39.7 (11.6) | 21 (4.9) | | | 1 (0) | 7.7 (0.7) | 0.67 (0.67) | ex. (right) |

Overview of electrode implantation and memory performance in Experiment 1.

^a^ Each number stands for the mean over all experimental sessions with the standard error across sessions in brackets.

^b^ SUs: Single Units (including ESNs)

^c^ ESNs: Episode Specific Neurons

^d^ SOZ: seizure onset zone; hip.: hippocampal; ex.: extrahippocampal; bi: bilaterally

Table S2.

Overview of electrode implantation and memory performance in Experiment 2.

| **Patient ID** | **Number of sessions** | **Number of bundles in hippocampus with neurons** | **Trial number**^a^ | | **Hits**^a^ | **Number of hipp. neurons^a,b^** | | **Number of ESNs^a,c^** | | **SOZ^d^** | |
| --- | --- | --- | --- | --- | --- | --- | --- | --- | --- | --- | --- |
| 1013 | 1 | 1 | 49 (0) | 37 (0) | | | 16 (0) | | 3 (0) | | ex. (right) |
| 1014 | 1 | 1 | 70 (0) | 28 (0) | | | 1 (0) | | 0 (0) | | hip (right) |
| 1015 | 3 | 1.7 | 62 (0.6) | 28 (1.8) | | | 9 (0.6) | | 2.3 (0.3) | | hip (right) |
| 1016 | 3 | 2 | 46.7 (4.1) | 31.3 (3.8) | | | 17.3 (1.5) | | 3.3 (0.3) | | ex. (bi) |
| 1017 | 2 | 1.5 | 62 (2) | 38.5 (0.5) | | | 2 (1) | | 0 (0) | | hip (right) |
| 1018 | 2 | 3 | 56 (9) | 31 (5) | | | 16 (1) | | 1.5 (0.5) | | ex. (right) |
| 1019 | 1 | 2 | 64 (0) | 49 (0) | | | 12 (0) | | 5 (0) | | ex. (left) |
| 1020 | 1 | 2 | 51 (0) | 27 (0) | | | 22 (0) | | 2 (0) | | ex. (right) |
| 1021 | 1 | 2 | 54 (0) | 38 (0) | | | 15 (0) | | 1 (0) | | ex. (left) |
| 1022 | 1 | 2 | 62 (0) | 28 (0) | | | 10 (0) | | 2 (0) | | hip (right) |
| 1024 | 1 | 1 | 78 (0) | 70 (0) | | | 3 (0) | | 1 (0) | | no seizures |
| 1026 | 1 | 2 | 49 (0) | 38 (0) | | | 15 (0) | | 3 (0) | | hip (left) |
| 1027 | 1 | 2 | 52 (0) | 34 (0) | | | 3 (0) | | 0 (0) | | ex. (left) |
| 1028 | 1 | 1 | 45 (0) | 43 (0) | | | 4 (0) | | 1 (0) | | hip (left) |

^a^ Each number stands for the mean over all experimental sessions with the standard error across sessions in brackets.

^b^ SUs: Single Units (including ESNs)

^c^ ESNs: Episode Specific Neurons

^d^ SOZ: seizure onset zone; hip.: hippocampal; ex.: extrahippocampal; bi: bilaterally
